# dCas allele sequestration (das-CRISPR): A Versatile New Method to Achieve Monoallelic Gene Editing in Mouse Embryos and in cell culture

**DOI:** 10.64898/2026.06.03.729891

**Authors:** Ghassan Yehia, Jian Pan, Laura Servinsky, Xuening Hong, Peter Romanienko

## Abstract

CRISPR-Cas9 technology is a powerful tool extensively used for genome editing in mouse and many other species. *Streptococcus pyogenes* Cas9 efficiently cuts both alleles in mouse zygotes leaving many edited embryos without a functional protein that might be needed to sustain development, to survive postnatally or to reproduce, thus complicating its overwhelmingly advantageous use in making gene modifications. About 25% of mouse genes are essential for embryonic development and another 7% are necessary for fertility, thus for these genes it is desirable to maintain a functional allele to establish viable lines from CRISPR-Cas9 edited mouse embryos. However, exclusive monoallelic editing is challenging to achieve with current CRISPR methods. Controlling the activity of Cas9 in genome editing is an ongoing research field focused on developing new methods to curtail its damage caused by excess of on-target and off-target editing. In this study we describe a novel and a simple method, we termed das-CRISPR, for dCas allele sequestration in combination with CRISPR system, that allows monoallelic editing of targeted allele in mouse and in cultured cell lines. This method incorporates the use of a nuclease deficient deadCas9 (dCas9) present at higher levels than an active Cas9, both complexed with the same single guide RNA (sgRNA) sequence. We showed the delivery of the two proteins as ribonucleoprotein complexes (RNP) into mouse zygotes leads to the generation of viable and fertile mice carrying lethal mutations in an essential gene. We found that greater amounts of dCas9 RNPs bind and protect a target site while the lower amount of functional Cas9 RNPs accessed the unoccupied target site resulting in higher frequency of monoallelic gene editing, compared to using just Cas9 alone. We also showed this method can mitigate and control the activity of Cas9 in mouse NIH3T3 cells in culture to achieve monoallelic editing. This method is a versatile approach to controlling excessive Cas9 activity on-target and off-target both *in vitro* and *in vivo*.

## Introduction

Advances in genome editing technology enable rapid and precise changes to the genomes of numerous model organisms^1^. Particularly, CRISPR-Cas9 is a powerful genome-editing tool both successfully and extensively used to manipulate genomes in many cell types and organisms, including mouse embryos. The Cas9 nuclease is directed to a specific genomic locus by a programmable single guide RNA (sgRNA) where it creates a DNA double-strand break (DSB)^2,3^. DSBs can be repaired by several DNA repair pathways including non-homologous end joining (NHEJ), an error-prone repair mechanism which generates small insertion/deletion (indel) mutations resulting in frameshifts and potential loss of gene function or by homology-directed repair (HDR) which is harnessed to introduce specific nucleotide substitutions provided by exogenous DNA donors often as single-stranded OligoDeoxyNucleotide (ssODN) ^4–6^.

In mouse, about 25% of gene knockouts are embryonically lethal and another 7% are necessary for fertility^7–9^. CRISPR-Cas9 ribonucleoprotein complexes (RNPs) delivered into mouse zygotes efficiently cut both alleles leaving many edited embryos without a functional protein^10^. Thus, the overwhelmingly advantageous use of CRISPR-Cas9 could be a drawback for editing essential genes like those necessary for embryonic development, cell survival or reproduction^7,11,12^. Embryonic lethality due to uncontrolled Cas9 targeting activity of essential genes in mouse zygotes can be a barrier to producing knockout founder (G0) mice and run counter to the goal of developing mouse models^11^. Thus, it is desirable to maintain a functional allele to allow for survival of G0 mice that have embryonic lethal mutations. Monoallelic-specific editing in mouse embryos can be challenging to accomplish because cleavage efficiency of constitutively active Cas9 nuclease is spatiotemporally uncontrollable^10^. Numerous strategies have been developed to control Cas9 activity via modifying the Cas9 protein or its programming sgRNA molecules ^13–19^, but such strategies are exploratory in nature, are time consuming to establish and not amenable for use in all genome editing schemes. Thus far, successful monoallelic editing approach to modify only one of the two copies of a gene, often requires CRISPR-Cas9 RNPs targeting gene variations, like exploiting single nucleotide polymorphisms between two alleles^20–22^. The major drawback to this approach is the lack of variations between coding genes in commonly used inbred mouse lines^23^.

To mitigate cleavage efficiency of active Cas9 nuclease and minimize the potentially deleterious effect caused by biallelic editing of essential genes, several methods have been deployed to preserve a functional wild-type allele in the developing mouse embryo, thus increasing the likelihood to generate live, genetically engineered G0 mice. One method relies on the zygote HDR mechanisms to repair both alleles with a mixture of two ssODNs one carrying silent CRISPR-Cas9-blocking mutations and the other carrying the desired nucleotide substitutions^5,24^. Although this is a simple strategy to adopt, relying on both alleles to be repaired by HDR using precisely each donor ssODN is further complicated through competition with the NHEJ repair pathway^25–27^. Two additional methods were reported, this time involving mouse embryo manipulation for the purpose of controlling the spatiotemporal activity of Cas9 endonuclease. The two-cell embryo microinjection method consists of microinjecting CRISPR reagents into one blastomere, therefore restricting the space of Cas9 into one blastomere while the other blastomere remains intact and plays a complementary role to generate viable chimeric G0 mice^28,29^. Microinjecting one blastomere of a 2-cell mouse embryo could be challenging and time consuming; moreover, biallelic lethal mutations at the cellular level, or mutations affecting cell growth and differentiation produced in one blastomere would result in an animal developing from only the unedited cells^28^.

The second method reported involves transplanting a Cas9 RNP-injected pronucleus into a fertilized zygote that had been depleted of the parental pronucleus to generate a reconstructed normal diploid zygote. This nuclear transfer was shown to control spatiotemporally the Cas9 activity through cytoplasmic dilution, thus allowing the resulting embryos to survive carrying lethal mutations in an essential gene along with a functional unedited wild-type allele^30^. Although these methods offer the potential to generate viable G0 mice and subsequently mouse lines, they are difficult to establish in the setting of a typical core facility. These different, cumbersome approaches highlight the definitive need to control Cas9 activity and accomplish monoallelic editing of essential genes in general and in mouse embryos in particular.

Here, we present a simple and effective method to promote monoallelic editing by sequestering and protecting one allele from being edited in mouse zygotes. This new method involves using a mixture of active Cas9 and a catalytically dead Cas9 (dCas9)^31^ both complexed with the same sgRNA but with reduced concentration of Cas9 relative to dCas9. We demonstrate the feasibility of the method by generating monoallelic editing of viable G0 mice carrying a functional, unedited wild-type copy of an essential gene, *Acvrl1* and in the commonly targeted Gt(ROSA)26Sor locus as a proof of concept. We show that simple reduction of Cas9 concentration is not a barrier to its ability to effectively cleave both alleles. Moreover, we demonstrate the versatility of this method to control Cas9 activity not only *in vivo* but also in cell culture, opening the way for possible cell therapeutic applications. We name the method das-CRISPR for <u>d</u>Cas <u>a</u>llele <u>s</u>equestration-CRISPR

## Results

### Available CRISPR-Cas9 methods are not efficient in producing viable mice with embryonic lethal mutations in *Acvrl1* gene

CRISPR-Cas9 based genome editing of genes essential during embryonic development was shown to be possible in some circumstances using 2-cell embryo microinjection^28,29^ or a mixture of two ssODNs one carrying silent mutations and the other carrying the desired nucleotide substitutions to mimic monoallelic editing^5,24^. However, in the case of *Acvrl1*, a gene essential during embryonic development, we found it challenging to introduce a stop codon prematurely in the coding sequence. Homozygous *Acvrl1* KO mice die at mid-gestation (∼10.5 dpc) with severe vascular abnormalities^32,33^, however, mice heterozygous for the *Acvrl1* KO mutation were normal and fertile. We used CRISPR-Cas9 and an ssODN for the purpose of inserting a specific stop codon into Acvrl1 (R478X). Several electroporation sessions using C57BL/6J mouse zygotes^34,35^ were conducted in an attempt to generate monoallelic mutants, including co-electroporation of two ssODNs; one encoding the R478X change and a second encoding a silent R478R change to rescue the developing embryos^24^. These combined sessions generated 1443 electroporated embryos that were transferred into the oviducts of 58 pseudopregnant females which produced only 6 live pups, representing 0.4% of all embryos transferred (Table I). Those born were found to carry indels across wild-type allele or the modified wild-type with codon substitution R478R, but none carried the R478X mutation. Moreover, we had conducted two microinjection sessions of single blastomere microinjection of 2-cell embryos. These produced 469 live 2-cell embryos, with one blastomere microinjected with Cas9 RNP and the donor ssODN, that were transferred into the oviducts of pseudopregnant females. We expected that the 2-cell embryo microinjection approach will generate live adult chimeric mice carrying the R478X lethal mutation. Although we were successful in generating 43 live founders, none carried the mutation. Moreover, only 6 mice out of the 43, representing 1.3% of all embryos transferred, were found to carry indels in the *Acvrl1* gene (Table I). These results clearly suggested that using conventional CRISPR-Cas9 approaches to edit *Acrvl1* gene are inefficient and failed to produce live mice with the R478X mutation. CRISPR-Cas9-mediated genome editing is highly efficient in that both alleles often contain indels^10^ which may be lethal for developing embryos. Therefore, it became clear that is essential to protect one allele from Cas9 action to maintain its functionality and ensure cell survival during development while leaving an unprotected allele, within the same zygote, to be targeted and edited by active Cas9. We reasoned that a dCas9, or in principle any catalytically dead RNA-programable endonuclease, could play that role by binding and protecting a cognate target DNA sequence from the action of an active RNA-guided endonuclease.

**Table I:** Summary of experiments using CRISPR-Cas9 conventional approaches to edit *Acrvl1* an essential gene. Number of live mice with indels was determined using endonuclease T7 assay. The percentage (%) corresponds to the number of live mice out of the total number of embryos transferred.\

| Experiment | Number of embryos transferred | Number of mice born | Number of live mice with desired mutation | Number of live mice with indels (%) |
| --- | --- | --- | --- | --- |
| 100% Cas9 Electroporation | 1443 | 6 | 0 | 6 (0.4%) |
| 100% Cas9 2-cell microinjection | 469 | 43 | 0 | 6 (1.3%) |

### dCas9 *in vitro* binds and blocks endonucleases from cutting target DNA

To assess the ability of dCas9 to effectively bind and protect DNA sequences from cutting by Cas9, we used an *in vitro* Cas9 digestion assay in combination with different ratios of dCas9 to compete for cutting the target site. The DNA template used is a 345bp PCR product amplified from Rosa26 locus containing an XbaI restriction enzyme site located within a CRISPR-Cas9 target site when complexed with the sgRNA C408^36^ and a CRISPR-Cas12a target site when complexed with the crRNA C592^37^ (Fig1A). The *in vitro* digestion assay was performed in reaction mixtures that contained 100ng of DNA, Cas9:dCas9 RNP mix with either 1:1 or 1:3 ratio. Endonucleases, XbaI or CasRNP complexes, with or without dCas9, were added at 0 time point to concomitantly compete for DNA binding, mimicking accessibility to genomic DNA in cell nucleus. DNA digestions were incubated at 37°C for up to 24h and the reactions were arrested at several time point as indicated in Figure1. Blocking Cas9 efficiencies by dCas9 are interpreted by the diminishing release of digested fragments and the persistence of the uncut DNA fragment. In Figure 1B, we showed that Cas9 alone, at the amount used, is capable of cleaving most of the PCR product within 30 min (lane 4) and proceeds further when digestion is extended to 60 min or 90 min (lane 5 and 6 respectively). Whereas the addition of dCas9 at 1:1 ratio to Cas9 digestion mix, completely stopped the cleavage of the product beyond the initial partial digestion (compare lane 7 to lane 2) even when digestion was extended to 2 hours incubation (lane 8-12). To test for the specificity of the blockage, dCas9 complexed with C473, a sgRNA targeting *Acvrl1*, did not interfere with Cas9 digestion (compare lane 5, 9 and 12, Fig.1B). We also tested whether dCas9 binding to the DNA can interfere with the digestion of XbaI enzyme by physically covering the overlapping restriction site^38^. As expected, XbaI restriction enzyme can completely digest the DNA in the presence of non-targeting dCas9 complexed to C473 (lane 13) whereas dCas9 complexed with C408 can block DNA cleavage by the restriction enzyme (lane14). Next, we checked whether increasing the amount of dCas9 could further block the cleavage of the DNA fragment. Cas9:dCas9 ratio of 1:1 and 1:3 was used in comparison to Cas9 alone 1:0 (Fig. 1C). When comparing the 2 ratios, the release of the cleaved fragments appeared to diminish by increasing the amount of the dCas9 (compare lanes 11, 12 to lanes 5 and 6); moreover, the inhibition of Cas9 cleavage is persisting up to 24 hours of incubation in presence of dCas9. dCas9 binding to the double strand DNA fragment appears stable *in vitro* in accordance with previous reports^39,40^. Finally, we wanted to confirm the capability of dCas9 to block the activity of other RNA guided endonucleases, like for example CRISPR-Cas12a. Rosa26 Cas9 cut site overlaps the crRNA sequence of Cas12a^37^ (Fig.1A). As previously reported, we showed that Cas12a alone, at the amount used, can efficiently cleaves the Rosa26 345bp PCR product (lane14-18, Fig.1D). However, the addition of dCas9 at 1:1 or 1:3 ratio to Cas12a digestion mix, completely prevented the cleavage of the PCR product even when digestion was extended to 24 hours incubation (lanes 2-6 and 8-12 Fig.1D).

**Figure 1:**
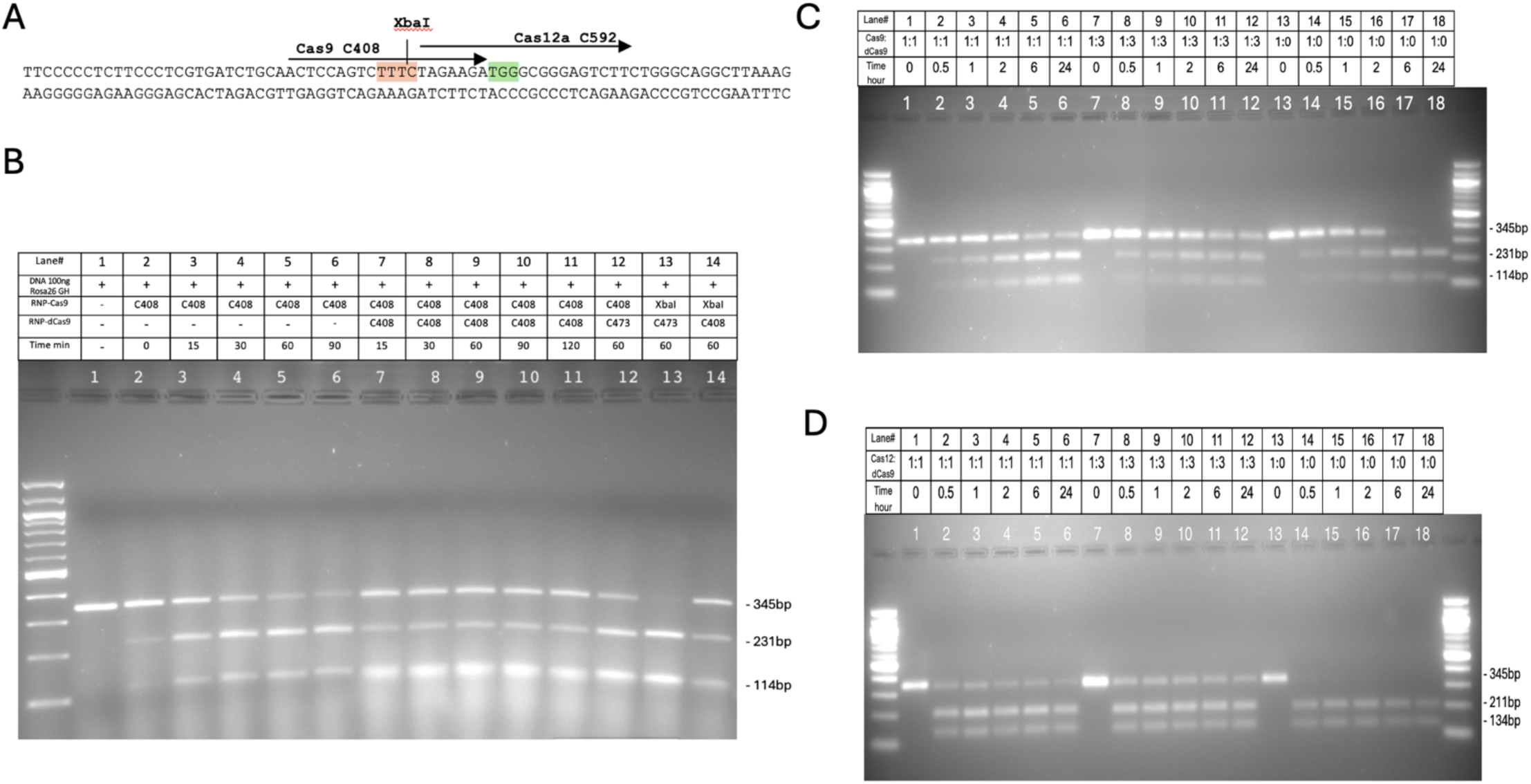
dCas9 *in vitro* modulate DNA digestion activity of three different endonucleases SpCas9, AsCas12a and XbaI. (**A**) partial DNA sequence of the purified 345bp PCR Rosa26-GH fragment, overlapping CRISPR- Cas9, -Cas12a and XbaI restriction enzyme cut sites. Protospacer sequences are highlighted and the PAM sequences are colored. Rosa26-GH fragment was subjected to *in vitro* digestion by three endonucleases XbaI and RNP complexes of Cas9 and Cas12a, with or without dCas9 RNP. DNA was added to the mix at 0 time point (no preincubation) and DNA digestions were let to incubate at 37°C. The reactions were arrested at several time point as indicated in the panel above each gel. Samples were run on 2% agarose gels stained with EtBr. 100bp DNA ladder used in each gel. (**B**) Cas9-RNP and dCas9-RNP associated with C408 sgRNA compete for the same site on the Rosa26-GH DNA fragment, while dCas9-RNP associated with C473 sgRNA is not specific to Rosa26. The ratio used of Cas9 to dCas9 was 1:1. XbaI restriction site is located 5bp downstream from Cas9 cut site. Digestion of Rosa26-GH fragment, by Cas9 or XbaI, releases two DNA fragments of approximately 231bp and 114bp. Lane1, control non-digested DNA. Note the presence of undigested sgRNA molecules on the bottom of the gel at 100bp mark. (**C**) same as in (B) except two different Cas9:dCas9 ratios were used 1:1 (lane1-6) and 1:3 (lane7-12); Cas9 alone 1:0, with no dCas9, used as digestion control (lane13-18). (**D**) same as (C) except instead of Cas9, Cas12a-RNP associated with crRNA C592 targeting Rosa26 was used. Cas12a digestion of the Rosa26-GH fragment releases two DNA fragments of approximately 211bp and 134bp.

Similarly, we tested the capability of dCas9 *in vitro* to bind and protect a DNA PCR fragment amplified from *Acvrl1* locus from cleavage by Cas9. When dCas9 and Cas9 RNP complexed to the same sgRNA C473 were used for *in vitro* digestion assay, we found that dCas9 prevented the cleavage of PCR product as described for the Rosa26 fragment (supplementary Fig.1). In conclusion, *in vitro* dCas9 can effectively bind to double strand DNA blocking access to the cut site of three tested active endonucleases: CRISPR-SpCas9, AsCas12a and XbaI restriction enzyme. We then proceeded to determine if dCas9 *in vivo* can play a similar role protecting genomic DNA regions from cleavage.

### High amount of dCas9 RNP can effectively block Cas9 activity *in vivo* at *Acvrl1* **locus**

dCas9 is catalytically inactivated Cas9 protein that retains the ability to bind CRISPR sgRNAs and strongly bind to DNA in a programmable manner^31^. In correlation with what we have shown *in vitro*, we hypothesized dCas9 could sequester and protect DNA from the action of Cas9. Contrary to an *in vitro* assay using purified DNA, DNA in the nucleus of an early-stage mouse embryo is dynamic and bound by nucleoproteins^41^, potentially leading to dCas9 displacement from its binding site ^42,43^. Therefore, titration of an optimal ratio of dCas9 to Cas9 was necessary to ensure both accessibility to and suitable protection of the target site. Five preparations of varied dCas9:Cas9 ratios were tested: 100:0 which is 100% dCas9 with no active Cas9, 80:20, 50:50, 20:80 and 0:100 which is 100% Cas9 with no dCas9 added. The six RNP mixes complexed to C473 sgRNA (targeting *Acvrl1*) were then electroporated each into one hundred mouse zygotes. Half of the electroporated embryos from each set were cultured *in vitro* for 4 days to the blastocyst stage. The remaining half were transferred into 3 pseudopregnant females per dCas9:Cas9 ratio and implanted embryos were collected at 10.5 dpc for analysis.

The fully expanded blastocysts were analyzed by PCR for the efficacy of dCas9 to sequester the *Acvrl1* target site protecting it from the action of Cas9; we reasoned that scoring for the retention of *Acvrl1* wild type sequences, presumably not cut while in the presence of active Cas9, should act as a reporter on dCas9 modulation of Cas9 activity. Therefore, DNA was extracted from 20 to 25 fully expanded blastocysts per each electroporation mix and assayed by PCR. Amplicons containing the Acvrl1 CRISPR-Cas9 target site were subjected to next-generation sequencing (NGS) for deep coverage^44^. The curated results of the in-depth analysis are shown as number of reads in Table II and the percentage of wild-type sequence reads is plotted in Figure 2A. As expected, blastocysts generated from zygotes electroporated with Cas9 alone (0:100) or with 20:80 (dCas9:Cas9) mix did not yield significant wild-type sequences as both copies of *Acvrl1* were cleaved as evidenced by indels at the cut site. Surprisingly, the 50:50 mix did not show any improvement to yield wild-type sequences, detected as 0.1% wild-type sequences in total reads (Table II, Fig.2A), contrary to what we have observed in the *in vitro* assay when using a 1:1 ratio. However, when analyzing the blastocysts electroporated with 80:20 (dCas9:Cas9) mix, the percentage of wild-type sequence reads increased to 62%, a strong indication that the addition of excess mount of dCas9 at the 80:20 ratio essentially protects alleles from CRISPR-Cas9 cleavage.

**Figure 2:**
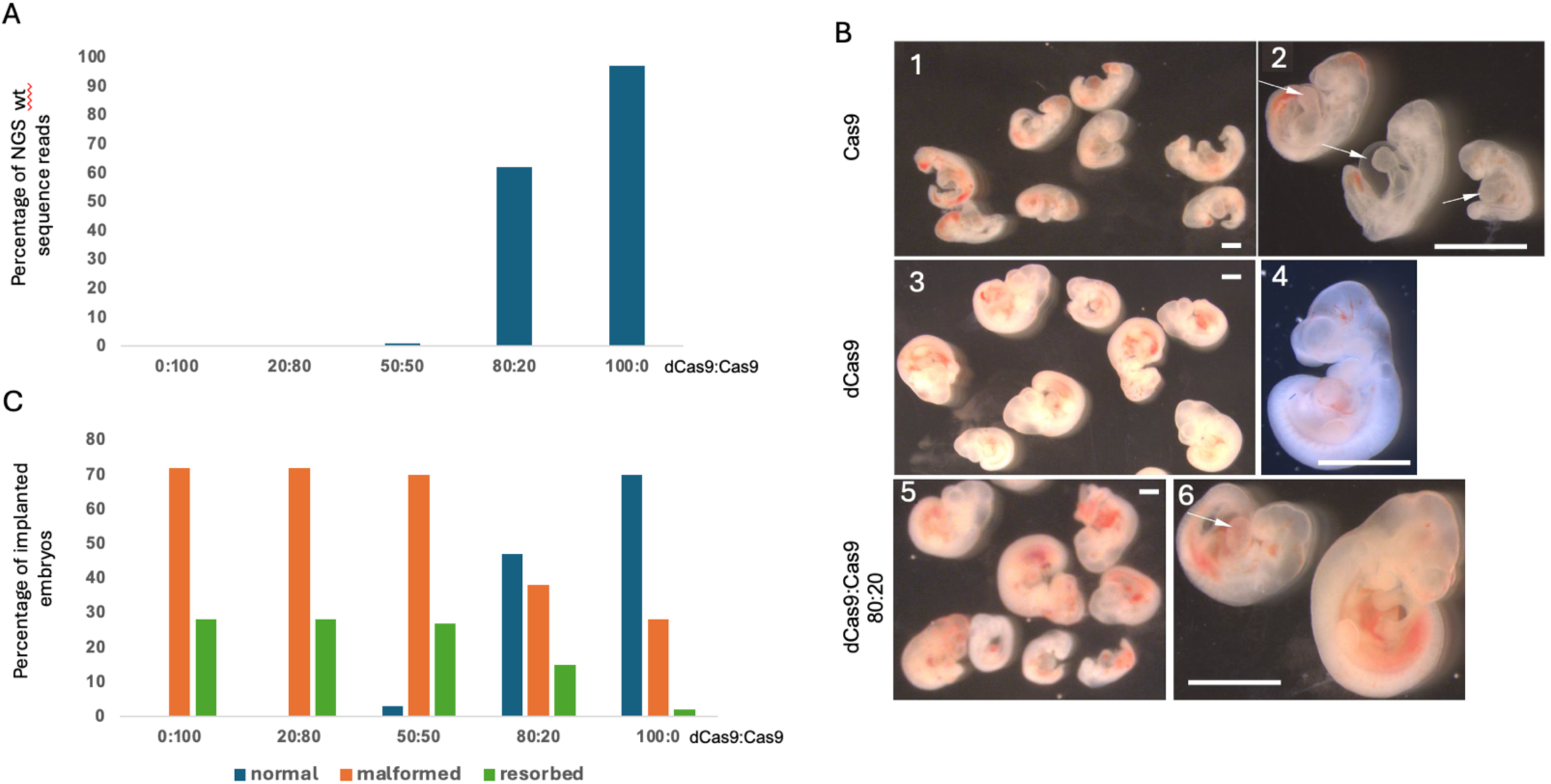
dCas9 *in vivo* blocks Cas9 activity from completely cleaving the *Acvrl1* target site and normal embryos are recovered. Five mixes of dCas9:Cas9 RNPs, associated with C473 sgRNA, of different ratios were electroporated into mouse zygotes. The ratio 0:100 is 100% Cas9 with no dCas9 added; on the other end 100:0 is 100% dCas9 with no active Cas9. (**A**) 50 electroporated zygotes from each condition were cultured *in vitro* to the blastocyst stage. 20-25 fully expanded blastocysts were collected per each ratio, pooled and lysed for genomic DNA extraction. Amplicons containing the *Acvrl1* CRISPR-Cas9 target site were sequenced by NGS and the curated results, are reported in Table II as total NGS sequence reads. The percentage of NGS wild-type reads from each ratio is plotted. This percentage, out of the total number of NGS reads, represent the proportion of uncut DNA fragments by Cas9. (**B**) 50 electroporated zygotes from each condition were transferred into three recipient females to develop *in utero*. After 10.5 dpc implanted embryos were collected, counted and examined; uterine resorption sites were also counted and included as implanted embryos, numbers are collected in Table III. Photomicrographs of E10.5 embryos collected from electroporation with Cas9-RNP (1, 2), with dCas9-RNP (3, 4) and with dCas9:Cas9 80:20 mix (5, 6). Embryos shown in (1, 3, 5) and in (2, 4, 6) were taken under same magnifications. (**C**) Embryos collected from each condition were classified as Normal-sized, Malformed and Resorbed embryos. The Y axis represent the number of classified embryos as a percentage of total number of counted embryos per ratio.

**Table II:**
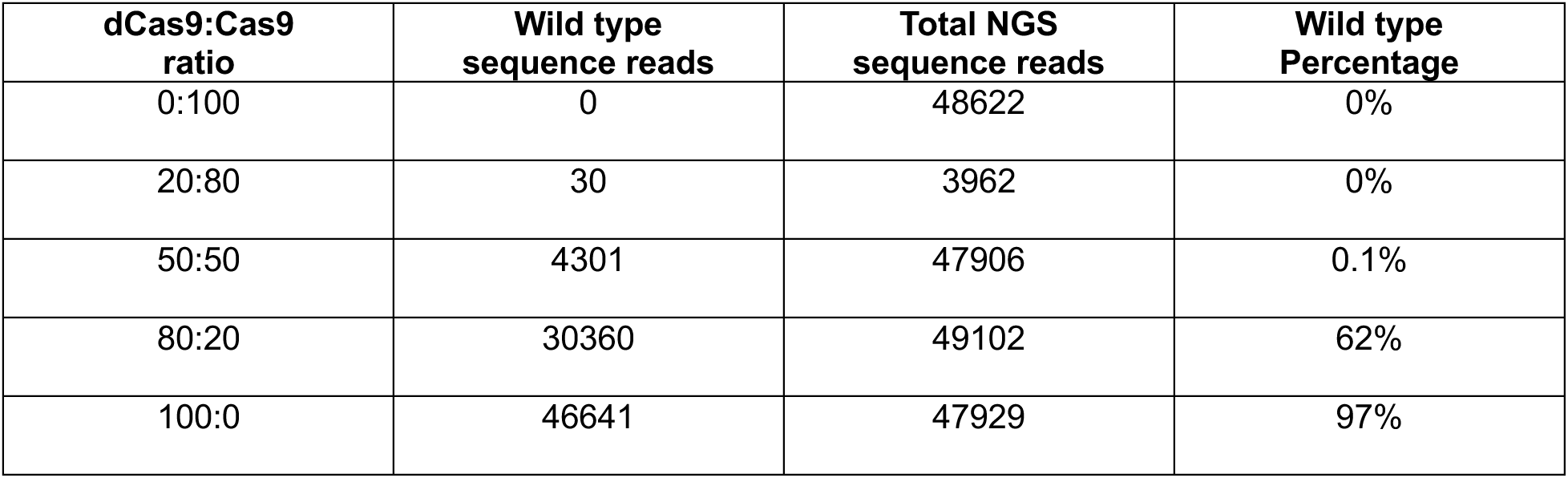
Summary of NGS reads from cultured blastocysts electroporated with five different ratios of dCas9:Cas9 RNP complexed with C473 sgRNA targeting *Acvrl1*. Genomic DNA extracted from 20-25 fully expanded blastocysts per ratio, used for PCR amplification of Acvrl1-AB fragment overlapping the Cas9 cut site. Amplicons containing the target site were sequenced by NGS and the results, curated by Azenta, reported in the table as total NGS sequence read. Wild-type percentage represents the number of all wild-type reads found out of the total number of NGS sequence reads.

**Table III:**
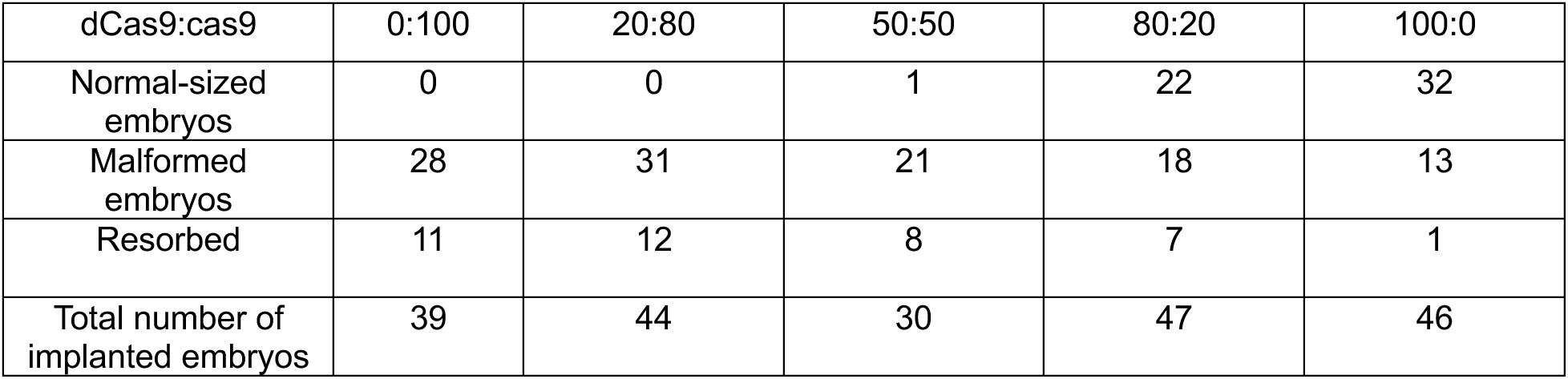
Summary of observation of embryos collected at 10.5 dpc from electroporation with different ratios of dCas9:Cas9 protein complexed with C473 sgRNA targeting *Acvrl1*. Observed embryos are classified as normal-sized, malformed or resorbed embryos. Numbers reported are the total number of embryos observed from 3 different uterine transfers.

Next, we investigated whether adding increasing amount of dCas9 would lead to increased number of embryo survival. The remaining electroporated zygotes with different ratios of dCas9:Cas9 were allowed to develop *in utero*. After 10.5 dpc, uterine horns were collected from pregnant females, resorption sites were counted and E10.5 embryos were isolated and their morphology examined. Homozygous *Acvrl1* KO E10.5 embryos had been reported to be distorted with severe growth retardation and enlarged pericardium^32,33^. As expected, all embryos electroporated with Cas9 only (0:100 ratio) showed morphology consistent with the published *Acvrl1* KO phenotype being reduced in size (Fig.2B1, 2B2) with an enlarged heart bud (arrows in Fig.2 B2). Whereas all embryos electroporated with dCas9 only (100:0 ratio) showed normal morphology with regards to embryo size and heart bud (Fig.2 B3, B4). However, when examining embryos from the electroporation of the 80:20 dCas9:Cas9 mix, 47% of embryos examined (Fig.2C) developed normally (Fig.2 B5, B6) with the remaining embryos showing reduced size and the characteristic enlarged heart bud (arrow in Fig.2 B6). Whereas all embryos collected from the 20:80 mix and all but one embryo from the 50:50 mix were found to be either malformed or resorbed (Table III, Fig.2C). This shows that the addition of excess dCas9 (80:20 ratio) is essential and sufficient for successful protection of an allele in a mouse zygote from CRISPR-Cas9 cleavage. It is noteworthy that the presence of normal 10.5 dpc embryos was reflected almost precisely by the percentage of wild-type sequence reads found in the cultured blastocysts (compare Fig.2A to Fig.2C). Thus, normal embryonic development could be rescued from the deleterious effect of uncontrollable active Cas9 by the addition of excess, which in this case was four-fold more, dCas9 protein.

**Figure 3:**
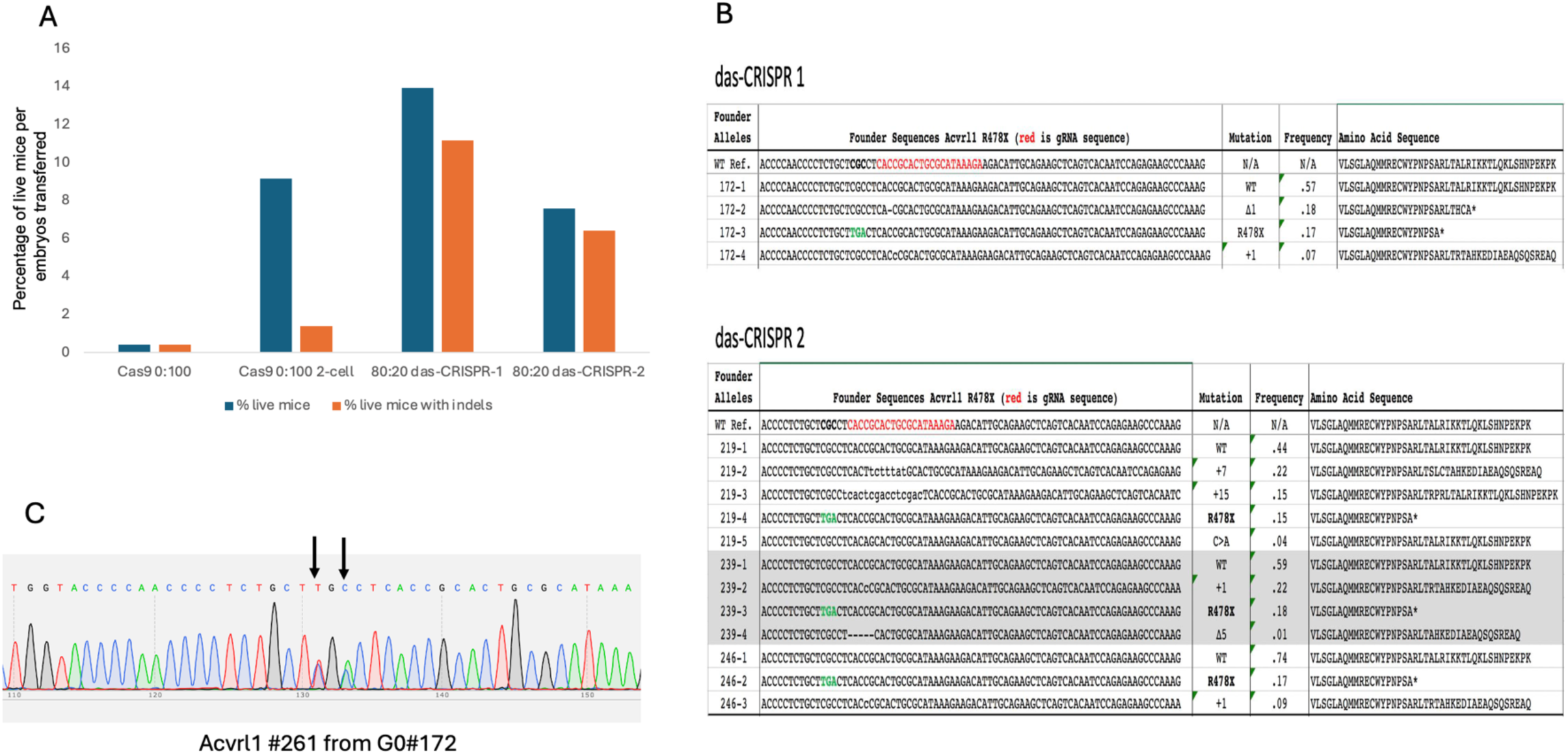
Editing of *Acvrl1* directly in mouse zygotes is possible when using das-CRISPR. (A) Live mice generated from either electroporated zygotes with 100% Cas9 (0:100) or microinjected of 2-cell embryos with 100% Cas9 (0:100) or electroporated zygotes with 80:20 ratio of dCas9:Cas9 mix from two sessions are plotted as percentage of live mice generated out of the total number of embryos transferred (numbers shown in table IV). (B) NGS reads of *Acvrl1* PCR fragment amplified from four live founders containing each, a wild-type allele in addition to alleles with indels and an allele with the R478X mutation. (C) Germline transmission of R478X allele to G1 generation. Representative sequencing chromatogram trace of ACVRL1-AB fragment from one descendant of G0 founder#172. Arrows shows the double peaks of base pair changes C/T and C/A confirming the codon change from CGC (Arg) to TGA (stop codon) in *Acvrl1*.

**Figure 4:**
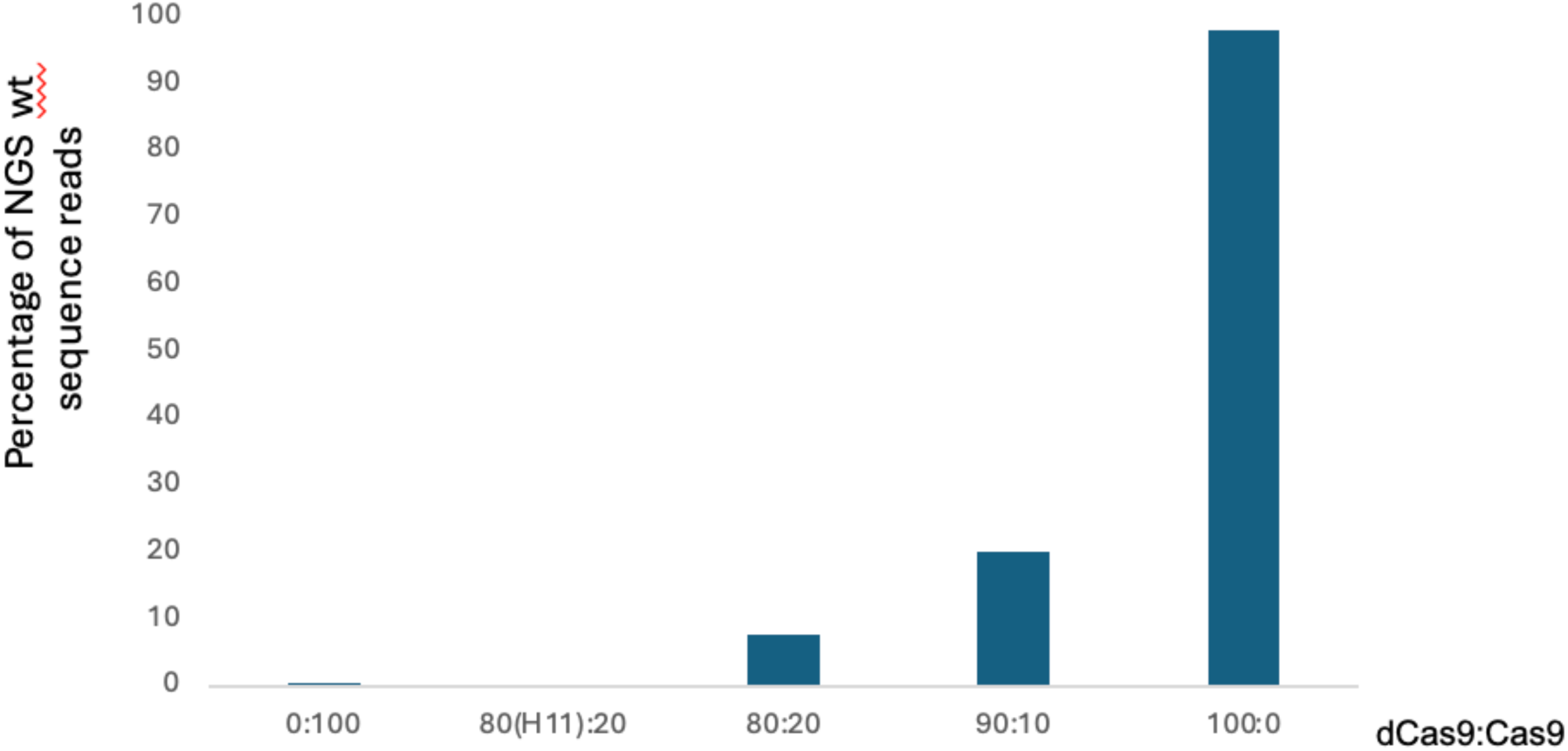
dCas9 targeting the Rosa26 *in vivo* specifically binds to the cut site and protect it from complete digestion by Cas9. Live mice were generated from zygotes electroporated with five different ratios of dCas9:Cas9 RNP complexed with C408 sgRNA targeting Rosa26 0:100, 80:20, 90:10,100:0, except for 80(H11):20 ratio where the dCas9 RNP was complexed to C433 sgRNA targeting the H11 genomic locus. Genomic DNA combined from 16 mice born per ratio is used for PCR amplification of Rosa26-GH fragment overlapping the Cas9 cut site. Amplicons containing the target site were sequenced by NGS and the results are reported as the percentage of wild-type reads out of the total number of NGS sequence reads.

**Figure 5:**
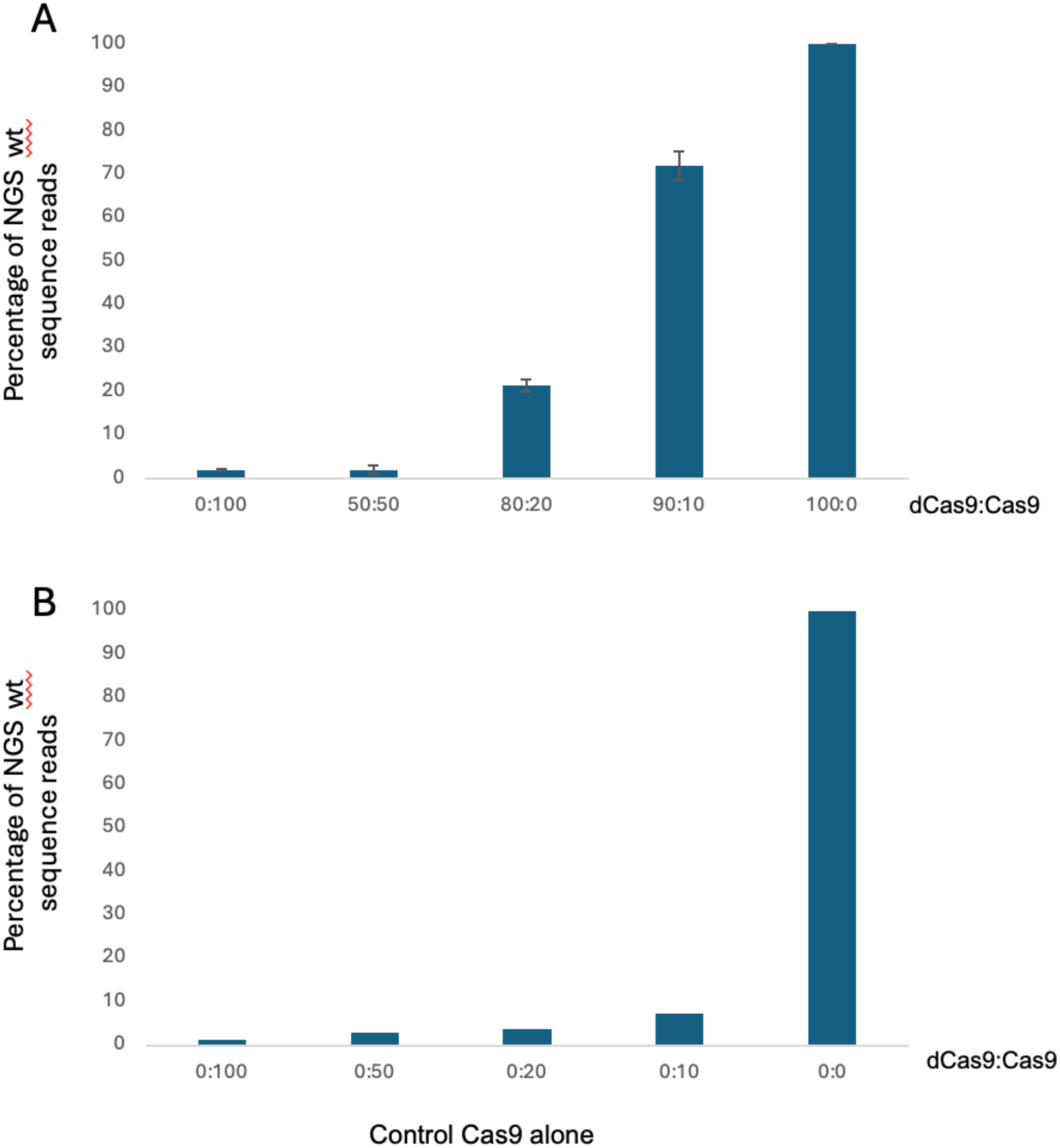
dCas9 binds and protects Rosa26 cut site from complete digestion by Cas9 in NIH3T3 cells in culture. NIH3T3 cells in culture were transfected with different ratios 0:100, 80:20, 90:10, 100:0 of dCas9:Cas9 RNP mix complexed with C408 sgRNA, targeting Rosa26. As a control, NIH3T3 cells were transfected with Cas9 RNP alone at the same concentration as above, except without dCas9, presented as 0:100, 0:50, 0:20, 0:10. The 0:0 ratio is a control without Cas9 or dCas9 RNPs in the transfection. Amplicons overlapping Cas9 cut site, from each ratio, were analyzed by NGS. The results of sequencing are plotted as percentage of wild-type reads out of the total sequence reads. The numbers plotted in (A) were the Mean values ±SD from two independent transfections.

**Figure 6:**
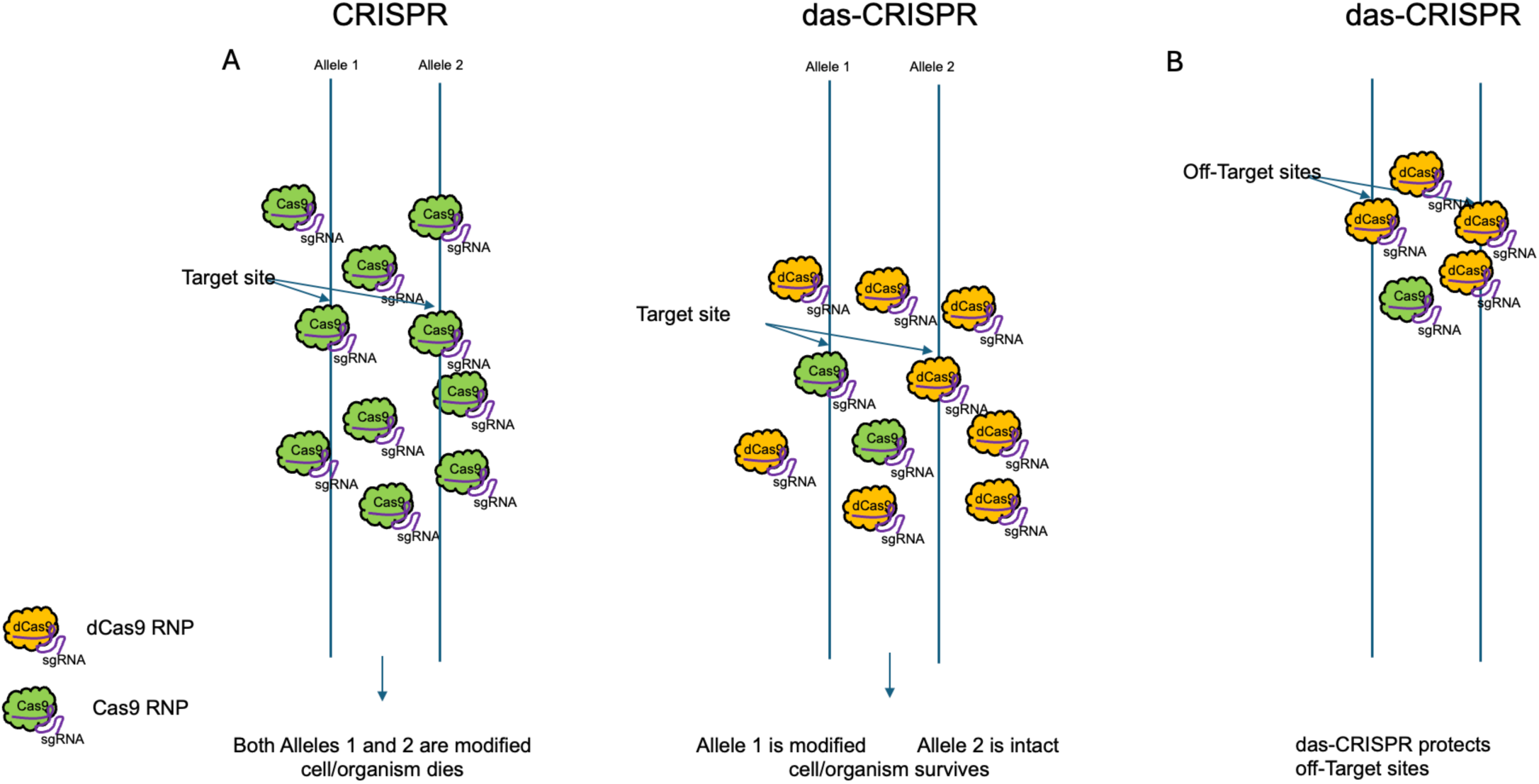
Schematic representations of das-CRISPR mode of action. (**A**) Right panel shows that das-CRISPR provides a means to selectively block one allele from modification, by utilizing dCas9 protein to compete for binding and thus protecting a target sequence from cleavage by a functional Cas9. Cas9 is then limited to target, when available, an allele unbound to dCas9 to induce a genetic change (right panel). If both alleles are inactivated (left panel) by Cas9, deleterious biallelic mutations can cause lethality. (**B**) das-CRISPR can be used to reduce off-target activity by directly competing with Cas9 for binding and protecting off target sites and thus increase editing efficiency. Cas9 and dCas9 RNPs, sharing the same affinity to off-target binding sites, are depicted competing for binding the same off target sites.

**Table IV:** Summary of experiments using CRISPR-Cas9 conventional approaches (from Table I) and from das-CRISPR to edit *Acrvl1* gene. The R478X desired mutation is screened by restriction digest. Number of live mice with indels was determined using endonuclease T7 assay. The percentage (%) corresponds to the number of mice out of the total number of embryos transferred.

| Experiment | Number of embryos transferred | Number of mice born (%) | Number of live mice with desired mutation | Number of live mice with indels (%) |
| --- | --- | --- | --- | --- |
| 100% Cas9 Electroporation | 1443 | 6 (0.4%) | 0 | 6 (0.4%) |
| 100% Cas9 2-cell microinjection | 469 | 43 (9%) | 0 | 6 (1.3%) |
| Das-CRISPR-1 80:20 | 215 | 30 (14%) | 1 | 21 (11%) |
| Das-CRISPR-2 80:20 | 528 | 40 (7.6%) | 3 | 34 (6.5%) |

### Successful generation of surviving mice with defined null mutation in *Acvrl1* when using das-CRISPR

Next, we wanted to ensure the rescued embryos were capable of developing beyond 10.5 dpc to produce live animals and that germline transmission of the Cas9-edited *Acvrl1* allele was possible. The dCas9:Cas9 ratio of 80:20 was chosen for proof of concept to finally generate live *Acvrl1* mutant mice either with a general null allele or with a defined null allele (R478X mutation) that could not be obtained using conventional CRISPR-Cas9 approaches. Two electroporation sessions were conducted: das-CRISPR-1 and das-CRISPR-2, each used dCas9:Cas9 80:20 RNP mix complexed with sgRNA C473, along with an ssODN containing the stop codon (R478X) as a donor template for HDR. Zygotes electroporated in each set were transferred into pseudopregnant females and 70 pups in total were born. It is noteworthy that these mice, to be able to survive, each must have a copy of a functional *Acvrl1* gene as heterozygous KO mice are reported to be normal^32^. This high number of born pups was in sharp contrast to previous attempts to generate *Acvrl1* mutants (results compounded in Table IV and in Fig.3A). From das-CRISPR-1, 24 pups (out of 30 born, representing 11% live mice out of total embryos transferred) and dAS-CRISPR-2, 34 pups (out of 40 born, representing 6.5% live mice out of total embryo transferred) were found to carry indels when analyzed by the T7 endonuclease assay (Fig.3A). Among the T7 positive mice, four were found harboring the desired R478X mutation when tested by restriction digest, corresponding to 0.52% live mice with desired mutation out of combined total embryos transferred. NGS analysis of the PCR amplicons containing the *Acvrl1* CRISPR-Cas9 target site, confirmed that those four mice were carrying *Acvrl1* alleles with the desired stop codon mutation, in addition to alleles with lethal indels and wild-type alleles found at higher frequency reads compared to the rest (Fig.3B). To test the transmissibility of lethal alleles, we crossed the four founders to C57BL/6J wild-type mice and genotyped G1 mice. The mutated alleles, including the desired R478X mutation, were found to be transmitted. Sanger sequencing trace from G1 mice carrying the R478X mutation showed the mutation change from CGC to TGA appearing as double peaks on the readout (see arrows in Fig.3C).

We demonstrated that the use of dCas9:Cas9 mix with 80:20 ratio can be used efficiently to generate mice with lethal mutations in an essential gene like *Acvrl1*. Subsequently, we wanted to explore what we expected to be the broad adaptability of this method to other genes.

### Targeting the Rosa26 locus with dCas9:Cas9 mix

Gt(ROSA)26Sor (Rosa26) is a non-essential gene used as a safe harbor for knock-in targeting using CRISPR-Cas9; moreover, CRISPR-Cas9 in combination with the sgRNA sequence of C408 was shown to exhibit high efficiency targeting Rosa26 in mouse zygotes^36^. Therefore, we wanted to test whether dCas9 would be able to moderate *in vivo* the high efficiency of Rosa26-targeted Cas9 RNP without bias towards embryo survival. We used for zygote electroporation of dCas9:Cas9 RNP mixes, complexed with C408 sgRNA, at different ratios 0:100, 80:20, 90:10 and 100:0 in addition to 80(H11):20 ratio where the dCas9 RNP was complexed to C433 sgRNA targeting the H11 genomic locus. H11 is unrelated yet similar to Rosa26 as a highly efficient CRISPR-targeted safe-harbor site^45^, which would not be expected to block Cas9 activity at Rosa26. These different ratio mixes were each electroporated into ∼ 100 zygotes, then transferred into pseudopregnant mice and allowed to develop to term. From each condition, 17-20 pups were born. Equal amount of genomic DNA from 16 born mice per condition were mixed and used to amplify by PCR the Rosa26 target site. Amplicons from each ratio were then analyzed by NGS for deep coverage of possible sequences generated. The results of sequencing are summarized as number of reads in Table V and plotted as percentage of wild-type reads (Fig.4). As expected, electroporation of Cas9 alone did not generate any significant number of wild-type reads whereas in combination with dCas9, 7.6% wild-type reads were generated when 80:20 ratio was used and 20% wild-type reads were seen with the higher ratio of 90:10. However, dCas9 RNP directed against H11 did not block Cas9 access to Rosa26, thus consequently no significant number of wild-type reads were generated (0.01%) confirming that the observed increase in wild-type reads is due to the protective effect of dCas9 binding and protecting the cut site from Cas9. Moreover, when the 80(H11):20 ratio was used, the concentration of 20% of Cas9 RNP was sufficient to generate indels at virtually all target sites just as effectively as 100% Cas9 RNP (Table V, Fig.4), therefore any effect of dCas9 on the observed increase of wild-type reads is due to its specific binding to genomic DNA and not to a reduced amount of Cas9 RNP in the ratio mixes. These results showed that dCas9 can be used *in vivo* efficiently to mitigate the effect of Cas9 in a non-biased development context and to preserve an intact copy of a targeted wild-type allele. As dCas9 functionally sequesters an allele from the cutting of Cas9, we termed the method das-CRISPR for dCas allele sequestration-CRISPR.

**Table V:** Summary of NGS reads from targeted Rosa26 locus. NGS reads from live mice generated from electroporated zygotes with five different ratios of dCas9:Cas9 RNP complexed with C408 sgRNA targeting Rosa26, except for 80(H11):20 ratio where the dCas9 RNP was complexed to a sgRNA targeting the H11 genomic locus. Genomic DNA combined from 16 mice born per ratio is used for PCR amplification of Rosa26-GH fragment overlapping the Cas9 cut site. Amplicons containing the target site were sequenced by NGS and the results, curated by Azenta, reported in the table as total NGS sequence read. Wild-type percentage represents the number of all the wild-type reads found out of the total number of NGS sequence reads.

| dCas9:Cas9 | Wild Type sequence reads | Total sequence reads | Wild Type Percentage |
| --- | --- | --- | --- |
| 0:100 | 436 | 122530 | 0.36% |
| 80(H11):20 | 14 | 104285 | 0.01% |
| 80:20 | 8236 | 107889 | 7.63% |
| 90:10 | 20908 | 104507 | 20.01% |
| 100:0 | 85213 | 86901 | 98.06% |

### dCas9 binds and protects Rosa26 cut site from complete digestion by Cas9 in cell culture

Next, we addressed the potential for das-CRISPR in cell culture as was similarly done *in vivo*. Cultured mouse NIH3T3 cells were transfected with dCas9:Cas9 RNP mixtures complexed with C408 sgRNA targeting Rosa26 locus, at these ratios: 0:100, 50:50, 80:20, 90:10 and 100:0. Each ratio mix was transfected using Lipofectamine into 50x10^5^ cells in 6-well plates in addition to controls with Cas9 RNP alone at different amounts corresponding to the different relative concentration of active Cas9 in the dCas9:Cas9 mixes, reported in Fig.5B as 0:100, 0:50, 0:20, 0:10. This was done to demonstrate relative cutting efficiency of limiting amounts of Cas9 in the absence of dCas9. We analyzed the outcome of each lipofection by PCR amplification of the target site followed by NGS deep sequencing (Fig.5). As for the *in vivo* experiments, we reasoned that scoring for the presence of Rosa26 wild-type sequences, which remained uncut, could act as a reporter on dCas9 modulation of Cas9 activity. Similar efficiency of cell transfection between wells was monitored by co-transfection of mCherry-expressing mRNA. Genomic DNA was extracted from each well after 2 days in culture and equal amount of DNA was used to amplify by PCR the Rosa26 target site. Amplicons from each ratio were then analyzed by NGS to detect all possible sequences generated. The results of sequencing are plotted as percentage of wild-type reads out of the total sequence reads (Fig.5). Low percentage reads of wild-type sequences are detected in cells transfected with Cas9 alone (0:100 ratio), in the experimental set and in the control set with 1.99±0.15% and 1.3% of total reads respectively; low reads of wild-type sequences remained unchanged, with the 50:50 ratio and the control 0:50 having 2.03±1.01% and 3% respectively (Fig.5A, B). However, the percentage of wild-type sequence reads increased to 21.4±1.36% for the 80:20 ratio and 72.06±3.3% for the 90:10 ratio (Fig.5A) which is a 10-fold and 35-fold increase, respectively, compared to Cas9 alone (1.99±0.15%). In the control set with Cas9 alone, but with lower concentration, the percentage of wild-type sequence reads remained low at 3.6% with 0:20 ratio and a slight increase to 7.4% with 0:10 ratio (Fig.5B). Therefore, the low representation of wild-type sequence reads observed in the control set with Cas9 alone, confirms that any effect of dCas9 on the increase of wild-type reads is due to its binding function to genomic DNA and not to a simple reduced amount of Cas9 RNP in any of the mixes. These results showed that dCas9 in cell culture can play a similar role as *in vivo* in modulating Cas9 activity and in protecting and preserving a wild-type copy of a functional allele.

## Material and Methods

### Animal experiments

All animal experiments were conducted on C57BL/6J mice (the Jackson Laboratory, Maine) in the Genome Editing Shared Resource of Rutgers-Cancer Institute of New Jersey, in accordance with U.S. National Institutes of Health (NIH) guidelines and regulations. All animal protocols were approved by the review board of the Institutional Animal Care and Use Committee of Rutger University. Mice were housed and maintained under specific pathogen-free conditions, in individually ventilated cages, under 12-hour light /12-hour dark cycle.

### Embryos electroporation and microinjection

Electroporation and microinjection were performed as described. Briefly, C57BL/6J zygotes collected from superovulated females, were either kept in M2 media for microinjection with Cas9 RNPs or rinsed through Opti-MEM media (Thermo Fisher Scientific) for electroporation and placed into total volume of 20μl 1:1 mixture of Opti-Mem and CRISPR-Cas RNPs between 1mm gap electrode on a glass slide (CUY501P1-1.5; NEPA GENE, Japan) connected to NEPA21 Type II electroporator. Only the poring pulse was used and was set to voltage: 30 V, pulse length: 3.0 msec, pulse interval: 100 msec, number of pulses: 5, decay rate: 0%, and polarity: + (NEPA21 electroporator, NEPA GENE). Embryos were then recovered, washed and cultured in equilibrated KSOM-AA (MR-121-D Sigma-Aldrich) until ready for embryo transfers into pseudopregnant females.

### Preparation of CRISPR-Cas RNP mixes with sgRNAs and ssODN

Cas9, dCas9 and Cas12a proteins (Alt-R® S.p. HiFi Cas9 Nuclease V3, Alt-R® S.p. dCas9 Protein V3, Alt-R® A.s. Cas12a Nuclease Ultra) and the Ultramer ssODN donors were purchased from Integrated DNA Technologies Inc. (IDT). All sgRNAs were purchased from MilliporeSigma and were HPLC purified. For mouse embryo electroporation, Cas9 and dCas9 RNP complexes were first prepared independently in 100μl total reaction containing 0.6μM of either protein equivalent to 100ng/μl with 3-fold molar excess of sgRNA at 1.8 μM, equivalent to 60ng/μl, with or without ssODN donor template added to final concentration of 200ng/μl. Then, different volumes from each RNP preparation were taken to reconstitute the 20:80, 50:50, 80:20, 90:10 ratios of dCas9/Cas9 RNP mixes; 10 μl of each ratio mixed with 10 μl of Opti-Mem used to electroporate 50 to 60 mouse embryos. For 2-cell microinjection, half the amount of Cas9 RNP and ssODN were used.

The target sequence of sgRNA C473 used for targeting *Acvrl1* was TCTTTATGCGCAGTGCGGTG<u>AGG</u> (PAM is underlined). The ssODN sequence, which contained homology arm and the R478X mutation, was 5’-GTCCTCTCCGGGCTGGCCCAGATGATGAGAGAGTGCTGGTACCCCAACCCCTCTG CTt<u>GaCTC</u>ACCGCACTGCGCATAAAGAAGACATTGCAGAAGCTCAGTCACAATCCAG

AGAAGCCCAAAGTG-3’, R478X sequence change is in lower case, the change also modified the PAM sequence to avoid recutting. The ssODN sequence, which contained the R478R silent mutation was 5’- GTCCTCTCCGGGCTGGCCCAGATGATGAGAGAGT

GCTGGTACCCCAACCCCTCTGCTC<u>GaCTC</u>ACCGCACTGCGCATAAAGAAGACATTG CAGAAGCTCAGTCACAATCCAGAGAAGCCCAAAG-3’. HinfI restriction site is underlined in both sequences. The target sequence of sgRNA C408 used for targeting *Rosa26* was ACTCCAGTCTTTCTAGAAGA<u>TGG</u> (PAM is underlined). For the preparation of 80(H11):20 dCas9/Cas9 ratio, the target sequence of sgRNA C433 targeting the H11 genomic locus was AACACTAGTGCACTTATCCT<u>GGG</u>.

### Genotyping and DNA analysis

DNA was extracted, using Hotshot^46^ method, from cultured blastocysts, 10.5 dpc fetuses or born mice and used for genotyping by PCR. *Acvrl1* alleles, with indels or with the R478X mutation, were amplified by PCR using primers ACVRL1A 5’-CTGCTATGTCTCCCGATCCTGAG-3’ and ACVRL1B 5’- CTCAGCTGTATTTTTGGCTGGATG-3’. Amplified PCR ACVRL1-AB fragments were either digested with HinfI restriction enzyme (New England Biolabs), introduced by R478X and PAM change, or subjected to T7 endonuclease assay (New England Biolabs) then sent for Sanger or NGS sequencing (Azenta Life Sciences). Rosa26 alleles, with or without indels, were amplified using primers ROSA26G 5’-AGTGTTGCAATACCTTTCTGGGAG-3’ and ROSA26H 5’-GGCGGATCACAAGCAATAATAACCTG-3’. Amplified PCR ROSA26-GH fragments were sent for NGS sequencing (Azenta Life Sciences). All NGS curated data were provided by Azenta.

### NIH 3T3 cell culture and lipofection

Murine fibroblast NIH 3T3 cells were maintained in Dulbecco’s Modified Eagle Medium (DMEM; Gibco 11995-065) supplemented with 10% fetal bovine serum (FBS) and 1% penicillin–streptomycin under standard culture conditions (37°C, 5% CO₂). Cells were passaged at ∼70–80% confluency using routine trypsinization. For transfection, cells were seeded to achieve ∼70% confluence at the time of DNA delivery. RNP mixes, prepared as above, introduced into cells using Lipofectamine 3000 (Thermo Fisher Scientific) according to the manufacturer’s instructions except P3000 reagent was omitted; mCherry mRNA (TriLink Biotechnologies) was used as control for transfection efficiency. Briefly, RNPs–lipid complexes were prepared in Opti-MEM and added to cells culture medium, next day the wells were washed, and cells were further incubated for 48 to 72 hours. Cells were lysed and genomic DNA was column purified (Qiagen) for genotyping and downstream analyses.

### CRISPR-Cas In vitro Digestion assay

RNP mixes were prepared independently in 20μl reaction containing either Cas9, Cas12a or dCas9 at 250ng/μl with 3-fold molar excess of sgRNA equivalent to 150ng/μl. Then, for the digestion assay, 1μl of each RNP preparation was added to 100ng of PCR fragments either Rosa26-GH fragment or ACVRL1-AB fragment (described above) with or without dCas9 RNP with equal volume for the 1:1 ratio or 3 volumes for the 1:3 ratio, in presence of 1X NEB Cas9 buffer (NEB) for a total volume of 20μl. RNase A was added for 5min at the end time of each reaction to degrade excess of sgRNA molecules, the reaction is completely stopped with an SDS buffer containing Proteinase K and incubated for 15 min at 55°C for the release of the digested fragments. The digested fragments were then separated in 2% agarose gels. The sequence of the Cas12a crRNA C592 targeting Rosa26 was <u>TTTC</u>TAGAAGATGGGCGGGAGTCTT (PAM sequence underlined).

## Discussion

### Das-CRISPR a versatile method that allows monoallelic editing in mouse zygotes

CRISPR technology is a very efficient tool which has been used to manipulate the genome in myriads of cell types and organisms including in mouse embryos. However, the CRISPR-Cas9 system has limitations, like uncontrolled activity of Cas9 leading to biallelic indels and potential off-target DSBs ^10,47–49^. About 25% of mouse genes are essential for embryonic development and perinatal survival with another 7% necessary for fertility and reproduction. CRISPR-Cas9 genome editing of essential genes in mouse zygotes, which could lead to embryonic or postnatal lethality, would be a consummate failure of the direct objective to generate G0 to perform long term studies.

Here we describe a new and simple method that uses a mix of dCas9 and Cas9 RNPs to achieve monoallelic editing of essential genes in mouse zygotes and efficiently generate adult mice carrying heritable lethal mutations. Dead Cas9 is a catalytically inactive Cas9 endonuclease harboring mutations in the RuvC1 and HNH nuclease domains (D10A and H841A) yet it retains its ability to bind specific DNA sequences when in association with sgRNAs^31^. The most common use of dCas9 widely reported as a tool for CRISPR-based gene regulation (CRISPRi/CRISPRa), imaging, and epigenetic modification ^50–52^. Here we demonstrate a new role for dCas9 to modulate Cas9 activity *in vivo* and to provide a new method for monoallelic editing. We report for the first time such utilization of dCas9 RNP to control the activity of Cas9.

We conducted several attempts using Cas9 alone in mouse zygotes to introduce a stop codon mutation into *Acvrl1* gene, an essential gene for embryonic development, however we failed to produce G0 mice carrying the intended R478X mutation (Table I, Fig.3A). This low number is consistent with what was previously reported about CRISPR editing of essential genes with embryonic or postnatally lethal knockout phenotypes ^8,11,12^. It noteworthy that the sgRNA sequence C407 used in these experiments, was very efficient in maximizing on-target cleavage of Cas9 RNP, as evidenced by the absence of wild-type NGS reads (Table II and in Fig.2A). Conversely, when we included an ssODN with a silent mutation (R478R) to rescue the developing embryos, we found low efficiency of donor-meditated HDR at the cut site, as only two pups out of the six born were found to carrying the R478R substitution.

Similarly, when active Cas9 alone was microinjected into one blastomere of 2-cell mouse embryos we failed to produce G0 mice carrying the R478X mutation, and more surprising out of the 43 mice we were able to produce, only six mice were found carrying lethal indels, representing 1.3% of live mice out of the total number of embryos transferred (Table I, Fig.3A). Extrapolating from the published method, we expected to find chimeric mice with lethal indels surviving as the descendants of both blastomeres in mouse zygotes contribute to the inner cell mass of blastocysts ^28^. We reasoned that Cas9 RNP efficiently induced biallelic lethal indels in the microinjected blastomere, most likely altering its function to properly contributing to the development of the resulting embryo leading to the generation of mice derived only from the uninjected blastomere.

Given the above challenges, it became evident that the use of only CRISPR-Cas9 to edit essential genes like *Acvr1* may not be practical and to achieve successful editing is conditioned on protecting a functional wild-type allele from the action of Cas9 endonuclease in the milieu of a dynamic, developing embryo. Much like a DNase protection assay^53^, we thought of using dCas9 to bind and protect the target site from the action of active Cas9, thus preserving a functional wild-type allele.

We first tested the capability of dCas9 to modulate Cas9 activity using an *in vitro* Cas9 digestion assay with a PCR product amplified from Rosa26 locus overlapping an XbaI cut site, a CRISPR-Cas9 cut site and a CRISPR-Cas12a cut site. The dCas9:Cas9 mixes were directly added, concomitantly competing for DNA binding, to assess the dynamic between the paired complexes. dCas9, when mixed with Cas9 at two different ratios, can modulate Cas9 endonuclease activity through competitive binding to the same target site and by sterically blocking the access to the cut site not only for Cas9 but also for Cas12a and XbaI restriction enzyme (Fig.1). Although Cas12a and Cas9 binding sites partially overlap (Fig.1), dCas9 footprint on the double strand DNA is reported to cover approximately 40bp, beyond the apparent interaction site of 20bp, which is enough to overlap Cas12a binding site and physically interfere with its function ^38,54^. Moreover, we showed that dCas9 modulation of Cas9 activity is sequence specific, dCas9 and Cas9 RNPs, complexed with the same sgRNA, physically compete to the same target site, whereas a dCas9 RNP complexed with an sgRNA sequence unrelated to the target site is uncapable of interfering with Cas9 endonuclease activity (Fig1 lane 12 and supplementary Fig1 lane 13). The next step was to extend these findings *in vivo*, i.e. modulating and interfering with Cas9 activity to prevent it from deleteriously cutting both alleles in mouse embryos.

Titration of an optimal ratio of dCas9 to Cas9 was necessary to ensure continuous accessibility and protection of the target site, the genome in the nucleus of early-stage mouse embryo is dynamic^41^, presumably leading to dCas9 displacement from its protective binding site^42,43^ thus requiring excess amounts of dCas9 to re-bind to target DNA and to persist after each blastomere division or DNA duplication until both RNPs are eventually depleted. The half-life of Cas9 RNP is reported to be approximately 24 hours^55^, long enough to persist through two embryonic cell division. Several increasing ratios of dCas9 to Cas9 RNPs, targeting *Acvr1*, were electroporated into mouse zygotes (Table II and Fig.2). Half of the electroporated zygotes were cultured *in vitro* for 4 days to the blastocyst stage and half transferred into pseudopregnant females. Two reporting measures were taken to assess the outcome of dCas9 inclusion into the mix: (1) measuring at the molecular level the retention of wild-type alleles, presumably not cut while in the presence of active Cas9, using deep next-generation sequencing of the target DNA; (2) assessment of whether embryonic development is rescued by the presence of functional wild-type alleles when successfully protected by dCas9. NGS sequencing of blastocyst generated from zygotes electroporated with either Cas9 alone or with 20:80 mix or with 50:50 mix did not yield significant amount of wild-type sequence reads; however, when the 80:20 mix was used the percentage of wild-type sequence reads spiked to 62%. We did not expect a spike increase rather a gradual increase in wild-type sequence reads correlating with an incremental amount of dCas9 in each mix. We reasoned that the spike was a strong indication that addition of high amount of dCas9 (80:20 ratio) is essential to successful protection of a wild-type allele from CRISPR-Cas9 cleavage in the dynamic environment of a nucleus in early-stage mouse embryo and active competition for binding sites. Future work with more mixes of dCas9:Cas9, between 50:50 and 80:20, might still be needed to fine-tune the gradual effect of dCas9 addition on Cas9 activity, also, there may be locus-specific ratios that would need to be determined empirically.

Next, we investigated whether adding increasing amount of dCas9 would lead to an increased number of embryo survival. The remaining electroporated zygotes, transferred into pseudopregnant females, were allowed to develop *in utero* and at 10.5 dpc. embryos were collected and their overall morphology was assessed. Embryos electroporated with Cas9 only (0:100 ratio) or with 20:80 mix or with 50:50 mix all showed morphology reminiscent of the published *Acvrl1* KO phenotype with reduced embryo size and enlarged heart bud^32,33^; only one embryo, from the 50:50 mix, out of 30 analyzed was found of normal size. The addition of dCas9 at these ratios did not seem to interfere with the activity of Cas9 to access the locus. However, when the 80:20 mix was used, about one-half of the embryos collected were found to be normal with properly developed structures. This result clearly demonstrated that the inclusion of dCas9 at higher ratio can rescue normal embryonic development from the deleterious effect of an uncontrolled active Cas9. Thus, the high percentage of wild-type sequence reads detected by NGS at 80:20 ratio in the cultured blastocysts is directly translated into greater embryo survival at 10.5 dpc (compare Fig.2B to Fig.2C). This is the first instance of dCas9 use *in vivo* to mitigate the activity of Cas9 and to potentially retain an intact allele during gene editing.

The question remained whether the rescued embryos at 10.5 dpc could fully develop to term and the pups could survive and transmit the KO or edited alleles. To determine if the normal E10.5 embryos survive postnatally, we repeated the experiment with electroporated zygotes using only the dCas9:Cas9 ratio of 80:20 complexed with sgRNA C473, along with a ssODN containing the stop codon R478X as a donor template for HDR. The electroporated zygotes, from two independent sessions, yielded seventy pups which strongly suggested the embryos analyzed at 10.5 dpc can survive and develop to term; moreover, for the mice to survive, they must carry a functional copy of *Acvrl1* gene. Approximately 80% of the pups carried indels when analyzed with the T7 endonuclease assay. Among those mice with indels, four mice were found carrying the R478X mutation across a wild-type allele at near 50% frequency indicating one allele had been sequestered from Cas9 and the alleles with lethal mutations including R478X allele could be rescued. The presence of wild-type at high frequency could be accounted to the fact that each cell in the developing mouse embryo must contains a functional wild-type allele, in accordance with we have shown that 2-cell embryos microinjection of one blastomere with active Cas9 resulted in generating mice mostly with wild-type alleles (Table I and Fig.3A), likely due to biallelic lethal mutations at the cellular level.

Four G0 founders with the R478X mutation were generated albeit at low frequency representing 0.52% live mice with desired mutation out of total embryos transferred. We attribute this low frequency to the lower efficiency of ssODN donor-meditated HDR previously observed at the cut site with a different ssODN carrying non-lethal R478R silent mutation. Having four founders would still be considered a successful project with the numbers of embryos that had been manipulated and with the number of pups produced. Finally, the G0 founders were bred, found to be fertile and capably transmitted to the next generation the R478X mutation (Fig.3C).

We showed unambiguously that our new method, das-CRISPR, can be used to edit *Acvr1*, an essential gene, directly in mouse zygotes by sequestering and protecting one wild-type allele from the action of Cas9. In principle, dCas9 inclusion to modulate Cas9 activity should work for other genes and with other genome editing nucleases.

We opted for the often used Rosa26 locus to test whether dCas9 can modulate a highly efficient on-target Cas9 activity and whether it can bind and preserve a copy of a wild-type allele without prejudice of function and necessity for survival. Live mice were generated from zygotes electroporated with different ratios of dCas9:Cas9 RNP complexed with a Rosa26 sgRNA. Compiled genomic DNA from each group of mice was used to generate NGS reads of all possible sequences. The collected data confirmed the observed increase in wild-type reads is due to the protective effect of dCas9 by specifically binding and protecting the common cut site from cleavage by Cas9.

Importantly, the reduced concentration of 20% Cas9 RNP, alone without dCas9, was sufficient to generate indels at virtually all target sites just as effectively as 100% Cas9 RNP (Table V, Fig.4), indicating that any effect of dCas9 on the increase percentage of wild-type reads is again due to its sequestration of target sites and not to a reduced amount of Cas9 RNP in the mixes.

This set of experiments showed das-CRISPR is a versatile method that could be applied to different genomic loci to modulate highly active Cas9 and to preserve unedited wild-type alleles even at an accessible target site like Rosa26.

Our novel das-CRISPR method is a well-established method in our genome editing facility that has been successfully applied to varied *in vivo* genome editing projects and because it is extremely powerful and successful, it is our method of choice to conduct monoallelic editing, particularly to quickly generate knockout animal models with lethal phenotypes using CRISPR-Cas9. Other facilities and laboratories have also adopted our method and reported successful outcomes^56,57^.

### Das-CRISPR a versatile method that allows monoallelic editing in cell culture

The success of using dCas9 for the purpose of protecting and preserving one allele while modifying the second allele opened the way for a new efficient method that allows monoallelic editing not only in embryos, but also in cell culture. As a proof of concept, we tested the versatility of das-CRISPR in cultured NIH3T3 cell line. In a similar way to *in vivo*, we chose the Rosa26 locus to test whether dCas9 can modulate Cas9 activity and whether it can bind and preserve a copy of a wild-type allele. Results were similar to what we observed *in vivo*, confirming that the inclusion of dCas9 at higher ratios could modulate and interfere with Cas9 activity in cultured cells. Although we did not sort cells within the 80:20 or 90:10 groups to determine if the increase in wild-type allele sequences are derived from individual heterozygous cells, the 21% of wild-type reads are unlikely to be all generated from a subset of wild-type that had been fully protected by dCas9, rather, they are likely derived from cells that are heterozygous with an intact wild-type allele and a Cas9-edited allele, whereas in the 90:10 ratio we could expect higher occurrence of wild-type unedited cell clones. In agreement with our initial findings, Skarnes *et al.* in a preprint manuscript^58^ successfully reported using dCas9 in variable ratios to modulate Cas9 activity in human induced Pluripotent Stem cells (hiPSCs) in order to increase the percentage of heterozygous clones carrying the desired mutations. Although their success rates varied between ratios, the optimal ratios reported were for the 1:1 and 1:1.5 dCas9:Cas9 mix, contrary to the ratio 80:20 we found in NIH3T3 cells. These differences could be attributed to the cell lines used, NIH3T3 are rapid-growing aneuploid fibroblast cells known for high proliferation rates^59^ while iPS cells are mostly diploid^60^ or could be attributed to increasing amount of total proteins between each dCas9:Cas9 ratio used by the authors compared to a constant amount of total proteins between ratios used in our experiments, or simply due to the delivery methods to the cells. These findings strongly indicate that das-CRISPR is amenable to monoallelic editing in cell culture, although ratio titration might be needed between different cell lines for optimum results.

### Relevance and applications

Various approaches and methods have been developed ^13–19^ to control Cas9 activity. However, their roles in precise genome editing have not been thoroughly explored and are time consuming to fine tune and to establish as a standard, broadly amenable go-to method for genome editing. In this study we show that das-CRISPR, with the inclusion of dCas9 at a higher ratio than an active Cas9, can be simply implemented for *in vivo* editing of essential genes to generate mouse models and for editing cell lines cultured *in vitro*. Moreover, it is reasonable to extrapolate the use of dCas9, or any other catalytically dead RNA-programable endonuclease, to control and modulate the activity of other CRISPR associated proteins like Cas12a. It can also be used against other genome editing systems such as zinc-finger nucleases or transcription activator-like effector nucleases (TALENs) and base editors. The mechanism of action of das-CRISPR, portrayed in Figure 6, can be adapted to various genomic editors for example by designing sgRNAs targeting the same, or overlapping, DNA binding sites. Moreover, assuming shared DNA binding affinity, for example, between dCas9 and Cas9 RNPs, dCas9 would be expected to outcompete Cas9 for binding to the same off-target sites (Fig.6B). Therefore das-CRISPR method could also be used to reduce unintended off-target DSBs. This later property was exploited by Christou *et al.* in a recent publication^56^ to facilitate a specific modification of an individual gene located within a closely related cluster of paralogs, supporting the expected beneficial effect of das-CRISPR method on reducing off-target DSBs.

Finally, developing cell-based therapies utilizing CRISPR–Cas9 is hindered by a prolonged Cas9 activity causing excessive on-target and off-target DNA damage triggering the p53-induced DNA damage response (DDR) pathway, particularly in human induced pluripotent stem cells, in hematopoietic stem cells and other cells ^15,61–63^ resulting in reduced proliferation and survival of edited clones. The inclusion of dCas9 in das-CRISPR method could help reduce the overall mutational burden caused by Cas9, thus facilitate the production and the survival of edited clones.

### Conclusion

Das-CRISPR is a simple method that deploys dCas9 in a novel way to modulate and to counter biallelic editing of a constitutively active Cas9 nuclease. dCas9 action results from competing against Cas9 for binding to the same DNA target site, thus preventing Cas9 from causing DSBs. However, higher ratios of dCas9 to Cas9 are necessary to achieve monoallelic editing *i.e.* protecting and preserving one allele while allowing modification on the second allele. This method can have wide application as a tool for monoallelic editing, for example, of essential genes *in vivo* not only in mice but other animals as well, and *in vitro* for cell line editing particularly for therapeutic applications that requires the preservation of an essential allele and precise editing on the corresponding one, all the while reducing the overall cellular mutational burden caused by an active Cas9 endonuclease.

## Authorship contribution

G.Y. Design, Investigation, Methodology, Formal analysis, Writing – review & editing. J.P. Methodology, Writing – review & editing. X.H Methodology, Writing – review & editing. L.S. Methodology, Writing – review & editing. P.R. Conceptualization, Design, Investigation, Methodology, Formal analysis, Writing – review & editing.

## References

1. Pacesa, M., Pelea, O. & Jinek, M. Past, present, and future of CRISPR genome editing technologies. Cell 187, 1076–1100 (2024).

2. Jinek, M. et al. A programmable dual-RNA-guided DNA endonuclease in adaptive bacterial immunity. Science 337, 816–821 (2012).

3. Ran, F. A. et al. Genome engineering using the CRISPR-Cas9 system. Nat. Protoc. 8, 2281–2308 (2013).

4. Hsu, P. D. et al. DNA targeting specificity of RNA-guided Cas9 nucleases. Nat. Biotechnol. 31, 827–832 (2013).

5. Paquet, D. et al. EYicient introduction of specific homozygous and heterozygous mutations using CRISPR/Cas9. Nature 533, 125–129 (2016).

6. Yoshimi, K. et al. ssODN-mediated knock-in with CRISPR-Cas for large genomic regions in zygotes. Nat. Commun. 7, 10431 (2016).

7. Cacheiro, P. & Smedley, D. Essential genes: a cross-species perspective. Mamm. Genome O7. J. Int. Mamm. Genome Soc. 34, 357–363 (2023).

8. Dickinson, M. E. et al. High-throughput discovery of novel developmental phenotypes. Nature 537, 508–514 (2016).

9. White, J. K. et al. Sanger Institute Mouse Genetics Project. Cell 154, 452–464 (2013).

10. Wang, H. et al. One-step generation of mice carrying mutations in multiple genes by CRISPR/Cas-mediated genome engineering. Cell 153, 910–918 (2013).

11. Elrick, H. et al. Impact of essential genes on the success of genome editing experiments generating 3313 new genetically engineered mouse lines. Sci Rep 14, 22626 (2024).

12. Peterson, K. A. & Murray, S. A. Progress towards completing the mutant mouse null resource. Mamm. Genome O7. J. Int. Mamm. Genome Soc. 33, 123–134 (2022).

13. Hynes, A. P. et al. Widespread anti-CRISPR proteins in virulent bacteriophages inhibit a range of Cas9 proteins. Nat. Commun. 9, 2919 (2018).

14. Kawamata, M., Suzuki, H. I., Kimura, R. & Suzuki, A. Optimization of Cas9 activity through the addition of cytosine extensions to single-guide RNAs. *Nat*. Biomed. Eng. 7, 672–691 (2023).

15. Khajanchi, N. et al. Controlling CRISPR-Cas9 genome editing in human cells using a molecular glue degrader. Mol. Ther. Nucleic Acids 36, 102640 (2025).

16. Maji, B. et al. A High-Throughput Platform to Identify Small-Molecule Inhibitors of CRISPR-Cas9. Cell 177, 1067–1079 (2019).

17. Oakes, B. L. et al. CRISPR-Cas9 Circular Permutants as Programmable ScaYolds for Genome Modification. Cell 176, 254–267 (2019).

18. Richter, F. et al. Switchable Cas9 Current Opinion in Biotechnology. vol. 48 119–126, (2017).

19. Zou, R. S., Liu, Y., Wu, B. & Ha, T. Cas9 deactivation with photocleavable guide RNAs. Mol. Cell 81, 1553–1565 (2021).

20. Cattaneo, C. et al. Allele-specific CRISPR-Cas9 editing of dominant epidermolysis bullosa simplex in human epidermal stem cells. Mol. Ther. J. Am. Soc. Gene Ther. 32, 372–383 (2024).

21. Polikarpova, A. V., et al. CRISPR/Cas9-generated mouse model with humanizing single-base substitution in the Gnao1 for safety studies of RNA therapeutics. Front. Genome Ed. 5, 1034720 (2023).

22. Wei, Y., Yue, T., Wang, Y. & Yang, Y. Fertile androgenetic mice generated by targeted epigenetic editing of imprinting control regions. Proc. Natl. Acad. Sci. U. S. A. 122, 2425307122 (2025).

23. Arslan, A. et al. Analysis of structural variation among inbred mouse strains. BMC Genomics 24, 97 (2023).

24. Lackner, M., Helmbrecht, N., Pääbo, S. & Riesenberg, S. Detection of unintended on-target eYects in CRISPR genome editing by DNA donors carrying diagnostic substitutions. Nucleic Acids Res. 51, (2023).

25. Bunton-Stasyshyn, R. K., Codner, G. F. & Teboul, L. Screening and validation of genome- edited animals. Lab Anim 56, 69–82 (2022).

26. Mianné, J. et al. Analysing the outcome of CRISPR-aided genome editing in embryos: Screening, genotyping and quality control. Methods 121, 68–76 (2017).

27. Ruis, B. L., Bielinsky, A. K. & Hendrickson, E. A. Gene editing and CRISPR-dependent homology-mediated end joining. Exp. Mol. Med. 57, 1409–1418 (2025).

28. Wang, G., Li, C., He, S. & Liu, Z. Mosaic CRISPR-stop enables rapid phenotyping of nonsense mutations in essential genes. Development 148, (2021).

29. Wu, Y. et al. Generating viable mice with heritable embryonically lethal mutations using the CRISPR-Cas9 system in two-cell embryos. Nat. Commun. 10, 2883 (2019).

30. Li, Y. et al. Precise allele-specific genome editing by spatiotemporal control of CRISPR- Cas9 via pronuclear transplantation. Nat. Commun. 11, 4593 (2020).

31. Qi, L. S. et al. Repurposing CRISPR as an RNA-guided platform for sequence-specific control of gene expression. Cell 152, 1173–1183 (2013).

32. Oh, S. P. et al. Activin receptor-like kinase 1 modulates transforming growth factor-beta 1 signaling in the regulation of angiogenesis. PNAS 97, 2626–2631 (2000).

33. Urness, L. D., Sorensen, L. K. & Li, D. Y. Arteriovenous malformations in mice lacking activin receptor-like kinase-1. Nat. Genet. 26, 328–331 (2000).

34. Qin, W. et al. EYicient CRISPR/Cas9-Mediated Genome Editing in Mice by Zygote Electroporation of Nuclease. Genetics 200, (2015).

35. Tröder, S. E. et al. An optimized electroporation approach for eYicient CRISPR/Cas9 genome editing in murine zygotes. PloS One 13, (2018).

36. Chu, V. T. et al. EYicient generation of Rosa26 knock-in mice using CRISPR/Cas9 in C57BL/6 zygotes. BMC Biotechnol. 16, 4 (2016).

37. Liu, Z. et al. ErCas12a CRISPR-MAD7 for Model Generation in Human Cells, Mice, and Rats. CRISPR J. 3, 97–108 (2020).

38. Josephs, E. A. et al. Structure and specificity of the RNA-guided endonuclease Cas9 during DNA interrogation, target binding and cleavage. Nucleic Acids Res. 43, 8924–8941 (2015).

39. Saifaldeen, M., Al-Ansari, D. E., Ramotar, D. & Aouida, M. Dead Cas9-sgRNA Complex Shelters Vulnerable DNA Restriction Enzyme Sites from Cleavage for Cloning Applications. CRISPR J. 4, 275–289 (2021).

40. Yang, W. et al. Detection of CRISPR-dCas9 on DNA with Solid-State Nanopores. Nano Lett. 18, 6469–6474 (2018).

41. Eid, A., Rodriguez-Terrones, D., Burton, A. & Torres-Padilla, M. E. SUV4-20 activity in the preimplantation mouse embryo controls timely replication. Genes Dev. 30, 2513–2526 (2016).

42. Brüning, J. G., Howard, J. A. L., Myka, K. K., Dillingham, M. S. & McGlynn, P. The 2B subdomain of Rep helicase links translocation along DNA with protein displacement. Nucleic Acids Res. 46, 8917–8925 (2018).

43. Kiernan, K. A. & Taylor, D. W. Visualization of a multi-turnover Cas9 after product release. Nat. Commun. 16, 5681 (2025).

44. Fernández, A., et al. Simple Protocol for Generating and Genotyping Genome-Edited Mice With CRISPR-Cas9 Reagents. (2020).

45. Browning, J. et al. Highly eYicient CRISPR-targeting of the murine Hipp11 intergenic region supports inducible human transgene expression. Mol. Biol. Rep. 47, 1491–1498 (2020).

46. Truett, G. E. et al. Preparation of PCR-Quality Mouse Genomic DNA with Hot Sodium Hydroxide and Tris (HotSHOT). BioTechniques 29, 52–54 (2000).

47. Chen, Q. et al. Genome-wide CRISPR oY-target prediction and optimization using RNA-DNA interaction fingerprints. Nat. Commun. 14, 7521 (2023).

48. Oliver, D., Yuan, S., McSwiggin, H. & Yan, W. Pervasive Genotypic Mosaicism in Founder Mice Derived from Genome Editing through Pronuclear Injection. PloS One 10, (2015).

49. Yen, S. T. et al. Somatic mosaicism and allele complexity induced by CRISPR/Cas9 RNA injections in mouse zygotes. Dev. Biol. 393, 3–9 (2014).

50. Gilbert, L. A. et al. Genome-Scale CRISPR-Mediated Control of Gene Repression and Activation. Cell 159, 647–661 (2014).

51. Maeder, M. L. et al. CRISPR RNA-guided activation of endogenous human genes. Nat. Methods 10, 977–979 (2013).

52. Perez-Pinera, P. et al. RNA-guided gene activation by CRISPR-Cas9-based transcription factors. Nat. Methods 10, 973–976 (2013).

53. Galas, D. J. & Schmitz, A. DNAase footprinting a simple method for the detection of protein-DNA binding specificity. Nucleic Acids Res. 5, 3157–3170 (1978).

54. Zhang, Q. et al. EYicient DNA interrogation of SpCas9 governed by its electrostatic interaction with DNA beyond the PAM and protospacer. Nucleic Acids Res. 49, 12433–12444 (2021).

55. Kim, S., Kim, D., Cho, S. W., Kim, J. & Kim, J. S. Highly eYicient RNA-guided genome editing in human cells via delivery of purified Cas9 ribonucleoproteins. Genome Res. 24, 1012–1019 (2014).

56. Christou, S. et al. Combining Cas9 and dCas9 facilitates genome editing in genes associated with viability or welfare issues, or within paralogous gene clusters. *bioRxiv* 2026.05.05.721005 (2026) doi:10.64898/2026.05.05.721005.

57. P-073: Use of deactivated Cas9 with active Cas9 to facilitate generation of mouse mutations associated with viability issues. in Abstracts of the 19th Transgenic Technology Meeting (TT2025 (eds Christou-Smith, S. et al.).

58. Skarnes, W. C. et al. Controlling homology-directed repair outcomes in human stem cells with dCas9. (2021).

59. Rahimi, A. M., Cai, M. & Hoyer-Fender, S. Heterogeneity of the NIH3T3 Fibroblast Cell Line. Cells 11, 2677 (2022).

60. Mayshar, Y. et al. Identification and classification of chromosomal aberrations in human induced pluripotent stem cells. Cell Stem Cell 7, 521–531 (2010).

61. Dorset, B., SR & R.O. The p53 challenge of hematopoietic stem cell gene editing. Mol Ther Methods Clin Dev (2023).

62. Haapaniemi, E., Botla, S., Persson, J., Schmierer, B. & Taipale, J. CRISPR-Cas9 genome editing induces a p53-mediated DNA damage response. Nat. Med. 24, 927–930 (2018).

63. Ihry, R. J. et al. p53 inhibits CRISPR-Cas9 engineering in human pluripotent stem cells. Nat. Med. 24, 939–946 (2018).

