## Supplemental Figure 1 for "dCas allele sequestration (das-CRISPR): A Versatile New Method to Achieve Monoallelic Gene Editing in Mouse Embryos and in cell culture"

**Supplementary Figure**

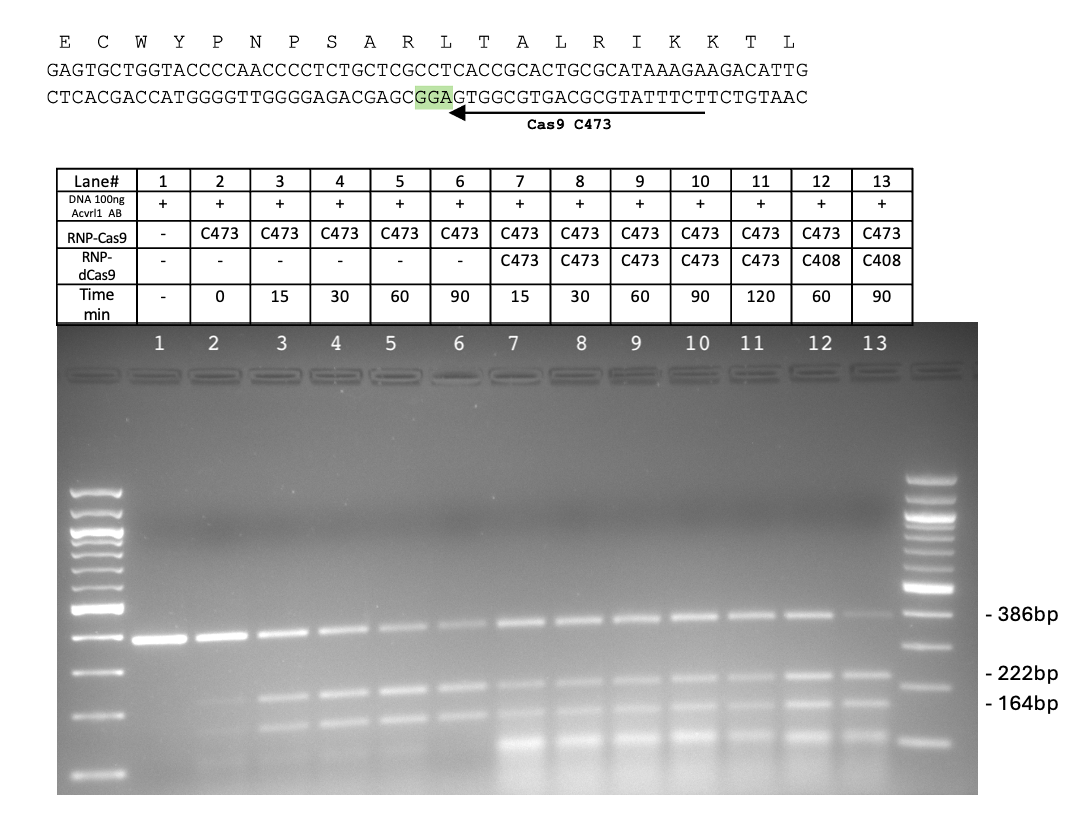

Supplementary Figure 1

**Supplementary Figure 1**: dCas9 *in vitro* blocks the digestion of purified PCR fragments, overlapping *Acvrl1* CRISPR-Cas9 cut site. Top: partial DNA sequence of the purified Acvrl1-AB 386bp PCR fragment, overlapping CRISPR-Cas9 cut site, the amino acid sequence is shown above it. Sequence of the protospacer C473 is highlighted and the PAM sequences is colored. Bottom: Acvrl1-AB fragment was subjected to *in vitro* digestion by Cas9 RNP with or without dCas9 RNP. DNA was added to the mix at 0 time point (no preincubation) and DNA digestions were let to incubate at 37°C. The reactions were arrested at several time point as indicated in the panel above the gel. Samples were run on 2% agarose gel stained with EtBr. 100bp DNA ladder used in each gel. Cas9-RNP and dCas9-RNP associated with C473 sgRNA compete for the same site on the DNA fragment, while dCas9-RNP associated with C408 sgRNA is not specific to *Acvrl1*. Cas9 digestion releases two DNA fragments of approximately 222bp and 164bp. Lane1, control non-digested DNA. Note the presence of undigested sgRNA molecules on the bottom of the gel at 100bp mark.
